# Oligomerization and cellular localization of SLC26A11

**DOI:** 10.1101/2024.04.29.591613

**Authors:** Stefanie Bungert-Plümke, Raul E. Guzman, Christoph Fahlke

## Abstract

The solute carrier family 26 (SLC26) encompasses multifunctional anion exchangers in all kingdoms of life. SLC26 proteins are known to assemble as dimers, and co-expression of multiple isoforms in certain cells raises the question whether different SLC26s can assemble into hetero-dimers. We focused on SLC26A11, a broadly expressed isoform that differs from other isoforms in its subcellular localization. Whereas the vast majority of SLC26-FP fusion proteins, i.e. SLC26A1, SLC26A2, SlC26A3, SLC26A4/pendrin, SLC26A5/prestin, SLC26A6, SlC26A7, and SLC26A9, localize to the surface membrane of transfected mammalian cells, we found exclusive lysosomal localization of SLC26A11. Renal collecting duct intercalated cells express SLC26A11 together with SLC26A4/pendrin and SLC26A7, and we therefore tested whether heterodimerization between these transporters might result in SLC26 transporter re-localization. Neither in HEK293T nor in immortalized intercalated cells co-expressing SLC26A11 with SLC26A4/pendrin or with SLC26A7, changes of SLC26A11 localization were observed. Moreover, native gel electrophoresis did not provide any evidence for heterodimerization of these isoforms. We next tested heterodimerization of SLC26A11 with SLC26A1, SLC26A2, SLC26A6 or SLC26A9 via co-expression in HEK293T cells and confocal imaging. For all combinations, no changes in subcellular distribution were observed. We conclude that SLC26A11 does not heterodimerize with other SLC26 proteins, and that heterodimerization does not target SLC26A11 to cellular surface membranes.

## Introduction

The SLC26 family contains ten human members that - with the exception of SLC26A5/prestin, the motor protein in outer hair cells (1) - function as transporters/channels for a broad range of anionic substrates (2,3). SLC26A1, SLC26A2, and SLC26A8 electroneutrally exchange SO_4_ ^2-^ or oxalate^2-^ with Cl^-^ or OH^-^. SLC26A3, SLC26A4/pendrin and SLC26A6 function as Cl^-^/HCO_3_^-^ exchangers. SLC26A7 (4,5) and SLC26A9 (6-8) have been postulated to function as anion channels. The physiological importance of SLC26-mediated transport proteins is highlighted by human diseases caused by mutations in SLC26 isoforms, such as dystrophic dysplasia for SLC26A2 (9), Pendred syndrome (PDS) for SLC26A4 (10), hearing impairment for SLC26A5/prestin (11), or elevated blood pressure for SLC26A7 (12).

SLC26A11 was cloned from guinea pig pancreatic duct (13) and simultaneously multiplied from an IMAGE clone after searching the translated human EST database for the closest human homologues of a yeast high affinity sulfate transporter (14). Initial characterization suggested SLC26A11 functioning as pH-dependent Cl^-^/SO_4_ ^2-^ exchanger. However, later reports assigned passive Cl^-^ currents in heterologous expression systems; SLC26A11 was therefore proposed to function as an anion channel (15) and hypothesized to regulate neuronal [Cl^-^] (16) or to mediate neuronal swelling in brain ischemia (17).

We found that heterologous expression of SLC26A11 results in preferential lysosomal localization, without discernible plasma membrane insertion. Since this localization is incompatible with the reported transport functions in cellular surface membranes, we tested whether SLC26A11 might reach the surface membrane of selected cells in a hetero-dimeric assembly with other SLC26s. Hetero-oligomerization often produces multimers with novel functional properties (18-21), but also modifies intracellular trafficking and thus targets subunits into cell organelles inaccessible to homooligomeric assemblies.

SLC26A11 co-exists together with SLC26A4/pendrin and SLC26A7 in intercalating cells (ICCs) of the renal collecting duct. Moreover, chondrocytes express SLC26A1, SLC26A2 and SLC26A11, or SLC26A11 might form heterodimers with SLC26A3, SLC26A6 or SLC26A9 in gastrointestinal and pulmonal epithelial cells. We tested whether these isoforms can assemble into heterodimers using heterologous expression in mammalian cells, confocal microscopy and biochemical approaches.

## Results

### SLC26A11 preferentially localizes to lysosomes in cultured cells

Figure 1 depicts representative confocal images from HEK293T cells expressing fluorescent fusion proteins of human (h)SLC26A1, hSLC26A2, hSLC26A3, hSLC26A4/pendrin, rat (r)SLC26A5/prestin, hSLC26A6, mouse (m)SLC26A7, WT hSLC26A9, a truncated hSLC26A9™ mutant that improves surface membrane insertion (22), or mSLC26A11. Whereas all other isoforms exhibit predominant surface membrane insertion, we observed exclusive intracellular localization of mSLC26A11 (Fig. 1), in contrast to earlier findings (23). Protein expression level, transfection rate and protein distribution were relatively similar across all analyzed SLC26 proteins, except for SLC26A3 and A9. SLC26A3 expressed only at low levels, and both proteins exhibited an additional intracellular protein localization. To test whether SLC26A11 localization might be dependent on the transfected cell line or on protein expression level we compared the subcellular distribution in four cell lines using multiple expression plasmids. No differences were obtained between experiments with HEK293T, COS1, CHO or HEK293 cells, using expression plasmids with different promotors that might produce separate expression levels (Suppl. Fig. S1). Moreover, we observed similar intracellular localizations for mouse and human SLC26A11 (Suppl. Fig. S2).

**Figure 1.**
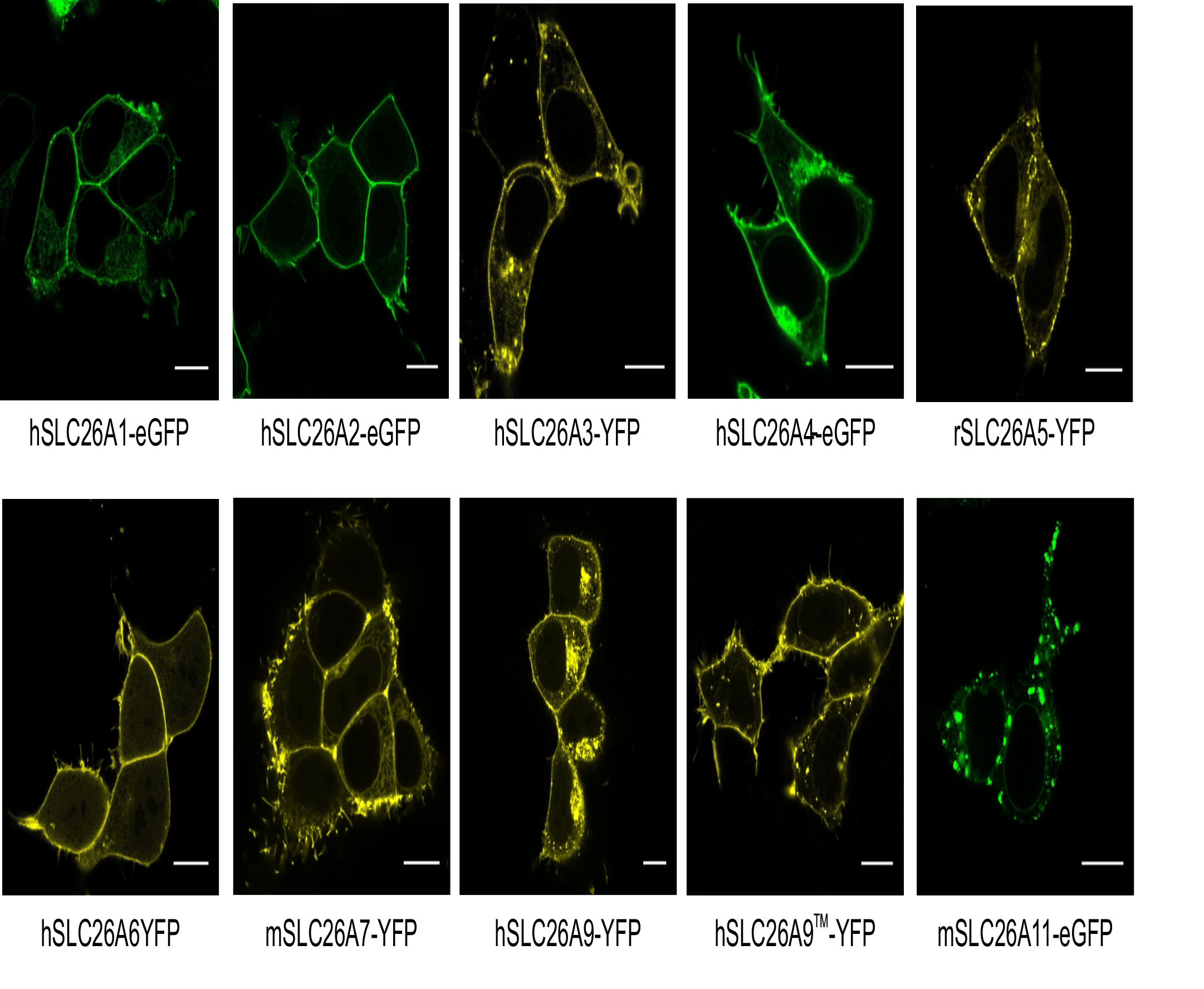
Expression of SLC26A proteins in HEK293T cells. Confocal images of HEK293T cells heterologously expressing SLC26A proteins tagged with indicated fluorescence proteins. All proteins are located in the plasma membrane, except for SLC26A11. Scale bar = 10 µm.

Intercalated cells in the mammalian collecting duct system express SLC26A4/pendrin, SLC26A7 and SLC26A11 (24). There exist two types of intercalated cells: α-intercalated cells (A-ICC) that absorb or ß-intercalated cells (B-ICC) that secrete HCO_3_^-^ (25). SLC26A11 was proposed to provide either an apical (in α-intercalated cells) or a basolateral (in ß-intercalated cells) Cl^-^ efflux pathway (24), and it is thus expected to insert into the surface membrane of these cells. We studied the cellular localization of SLC26A11 after heterologous expression in an immortalized bicarbonate secreting intercalated cell line, in Clone C cells (26). When seeded at low density, clone C cells differentiate into ß-intercalated cells with basolateral H^+^-ATPase and Cl^-^/HCO3^-^ exchanger AE, and into α-intercalated cells after seeding cells at high density (27,28).

Figure 2 illustrates the localization of SLC26A11 expressed either in HEK293T cells or in Clone C cells, either co-expressed together with Lamp1, a lysosomal marker protein, or with Calnexin, an endoplasmic reticulum (ER) marker. For both cell lines, the subcellular distribution of SLC26A11 significantly overlaps with the lysosomal marker protein Lamp1 (Fig. 2*A*), as judged by Manders coefficients ((MC)_HEK293T_ = 0.45 ± 0.11 (n=20) and (MC)_CloneC_ = 0.50 ± 0.09 (n=20), (Fig. 2*G*)). In contrast, we found only a small fraction of the SLC26A11 fluorescent signal overlapping with the ER marker Calnexin (MC-SLC26A11_HEK293T/CloneC_ = 0.12/0.09 ± 0.05/0.06 (n=19/18), (Fig. 2*B* and 2*H*).

**Figure 2.**
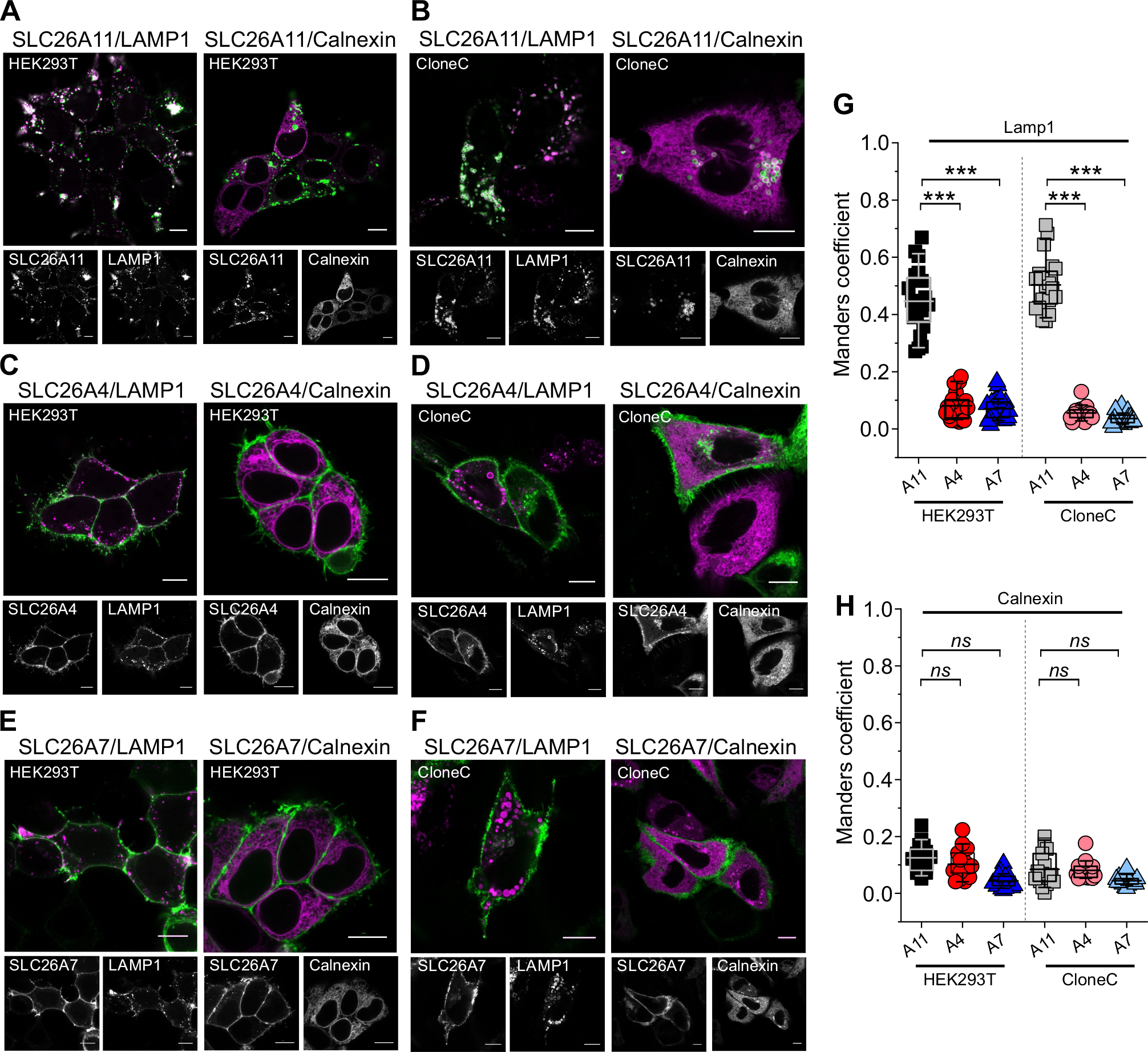
Subcellular localization of heterologously expressed SLC26A4/pendrin, SLC26A7 and SLC26A11. *A – F*, Confocal images of HEK293T cells (*A, C* and *E*) and B-ICC Clone C cells (*B, D* and *F*) co-expressing SLC26A4/pendrin, SLC26A7 and SLC26A11 together with the lysosomal marker Lamp1 or the ER marker Calnexin. *G* and *H*, box plot of Manders coefficients of colocalization analysis between SLC26A4/pendrin, SLC26A7 and SLC26A11 with the corresponding intracellular marker (SLC26A11/Lamp1_HEK293T_, n = 20; SLC26A11/Lamp1_CloneC_, n = 20; SLC26A11/Calnexin_HEK293T_, n = 19; SLC26A11/Calnexin_CloneC_, n = 18); (SLC26A4/Lamp1_HEK293T_, n = 20; SLC26A4/Lamp1_CloneC_, n = 14; SLC26A4/Calnexin_HEK293T_, n = 18; SLC26A4/Calnexin_CloneC_, n = 15); (SLC26A7/Lamp1_HEK293T_, n = 25; SLC26A7/Lamp1_CloneC_, n = 23; SLC26A7/Calnexin_HEK293T_, n = 25; SLC26A7/Calnexin_CloneC_, n = 21); One-way ANOVA (Tukey’s HSD post hoc test). Data are presented as means ± S.D. with n being the number of co-transfected cells analyzed. In boxplots, boxes indicate the upper and lower quartiles, and whiskers the upper and lower 90 percentiles. Scale bars = 10 µm.

SLC26A4/pendrin and SLC26A7 are predominantly located in the plasma membrane (Figs. 2*C*-2*F*) and cannot be detected together with Lamp1 or Calnexin (Figs. 2*C*-2*H*), (Lamp1, MC-SLC26A4_HEK293T/CloneC_ = 0.08/0.05 ± 0.05/0.03, (n=20/14); MC-SLC26A7_HEK293T/ CloneC_ = 0.07/0.04 ± 0.04/0.02 (n=25/23); Calnexin, (MC-SLC26A4_HEK293T/CloneC_ = 0.10/0.08 ± 0.05/0.03 (n=18/15) and SLC26A7_HEK293T/CloneC_ = 0.04/0.04 ± 0.02/0.01 (n=25/21).

### Lysosomal localization of SLC26A11-eGFP is not caused by the fluorescent protein tag

SLC26A11 was expressed as fluorescent fusion protein (SLC26A11-eGFP) in Figures 1 and 2 to facilitate visualization in confocal images. Since the added eGFP moiety might have modified the intracellular SLC26A11 localization, we compared untagged SLC26A11 with SLC26A11-eGFP in HEK293T cells and Clone C cells seeded at low (and therefore differentiated into B-ICC) or at high densities (and therefore differentiated into A-ICC) (Fig. 3). In these experiments, cells were fixed and stained with a newly generated anti-SLC26A11 monoclonal antibody (17D1). Detection with anti-rat Cy3 (for SLC26A11-eGFP (Figs. 3*A* and *B*)) or anti-rat Alexa488 (for untagged SLC26A11 (Figs. 3*C* and *D*)) revealed lysosomal localization of both tagged and untagged transporters. In both cell types and for every tested cell density, perfect overlap of eGFP with Cy3 fluorescence demonstrates that the 17D1 antibody detects SLC26A11 protein (Figs 3*A* and *B*). SLC26A11 also localizes to the lysosome with eGFP tag.

**Figure 3.**
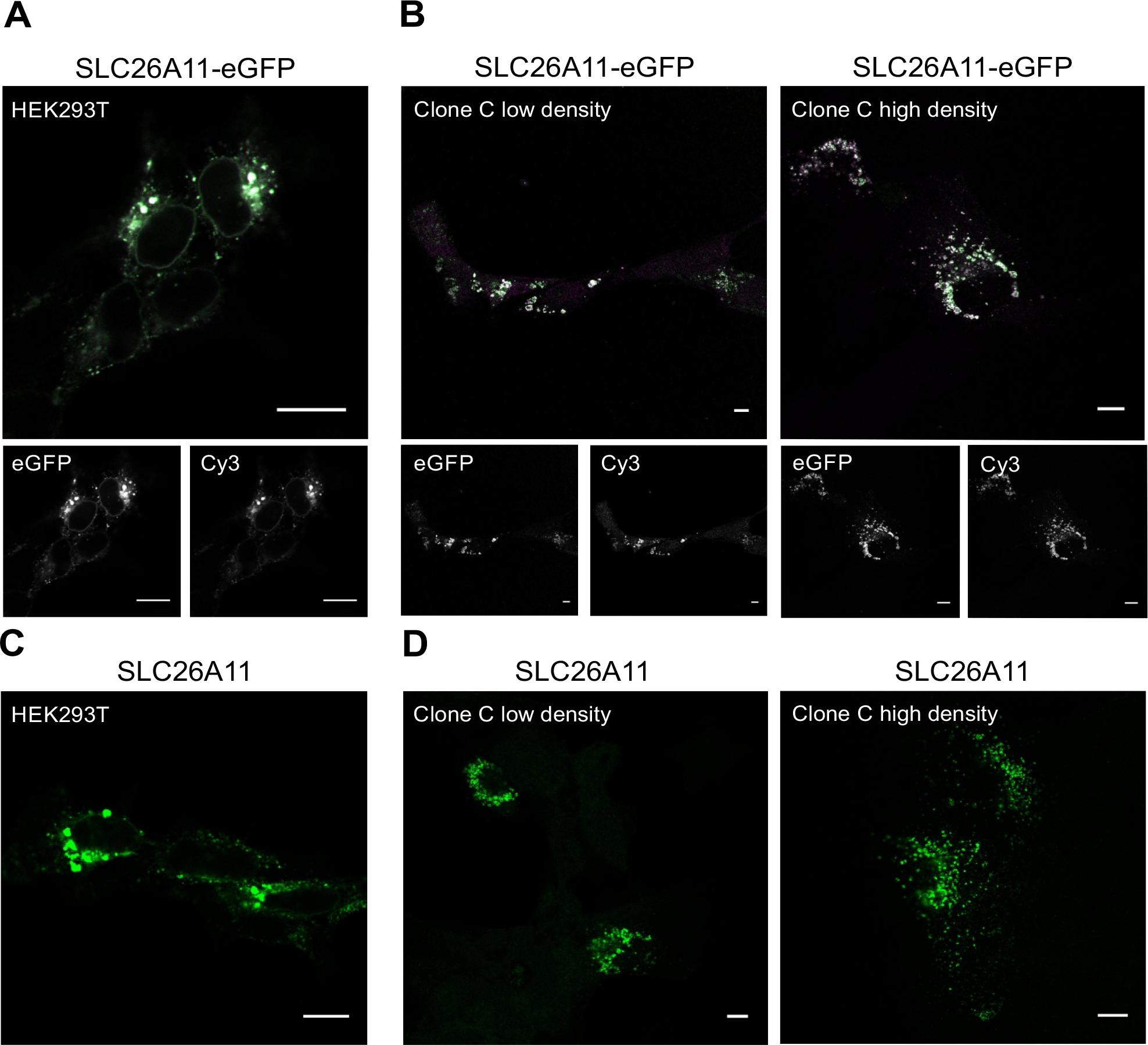
Detection of SLC26A11 with the anti-SLC26A11 monoclonal rt17D1 antibody. *A* and *B*, Confocal images of HEK293T cells (A), ß-intercalated Clone C cells (*B*, low dencity) or α-intercalated cells (*B*, high density) expressing SLC26A11-eGFP tagged protein. SLC26A11 was visualized directly via eGFP fluorescence or detected with the anti-SLC26A11 rt17D1 monoclonal antibody and visualized with donkey-anti-rat Cy3. *C* and *D*, Confocal images of both cell lines expressing untagged SLC26A11 protein. SLC26A11 was detected and visualized with anti-SLC26A11 rt17D1 antibody and donkey-anti-rat Alexa488. Scale bars = 10 µm.

### Co-expression with SLC26A4/pendrin or SLC26A7 leaves the subcellular distribution of SLC26A11 unaffected

Cl^-^ transport through the apical or basolateral membrane is believed to be a specific function of SLC26A11 in intercalated cells. Such a function is in disagreement with the lysosomal localization of SLC26A11 in transfected Clone C cells. Since SLC26A11 is co-expressed with SLC26A4/pendrin and SLC26A7 in these cells (24), and since hetero-oligomerization can target proteins into cell organelles inaccessible to homodimeric assemblies (29), SLC26A11 might have reached the surface membrane in a heterodimeric assembly. We therefore tested the subcellular localization of SLC26 isoforms in co-transfected HEK293T or Clone C cells.

Neither SLC26A4/pendrin nor SLC26A7 were able to change the subcellular localization of SLC26A11 (Fig. 4). The subcellular distributions of SLC26A4/pendrin and SLC26A11 barely overlap in both analyzed cell lines (Manders coefficient (MC)_HEK293T_ = 0.23 ± 0.1 (n=18) and Manders coefficient (MC)_CloneC_ = 0.24 ± 0.09 (n=13). SLC26A4/pendrin remained in the plasma membrane and SLC26A11 mainly in intracellular structures. Co-expression experiments of SLC26A11 together with SLC26A7 give even more pronounced results (Manders coefficient (MC)_HEK293T_ = 0.15 ± 0.06 (n=20) and Manders coefficient (MC)_CloneC_ = 0.05 ± 0.04 (n=25) with SLC26A7 being located in the plasma membrane and SLC26A11 intracellularly (Fig. 4*C*). We conclude that SLC26A11 does not heterodimerize with other SLC26A members such as SLC26A4/pendrin and SLC26A7 and that their co-expression does not modify their subcellular distribution.

**Figure 4.**
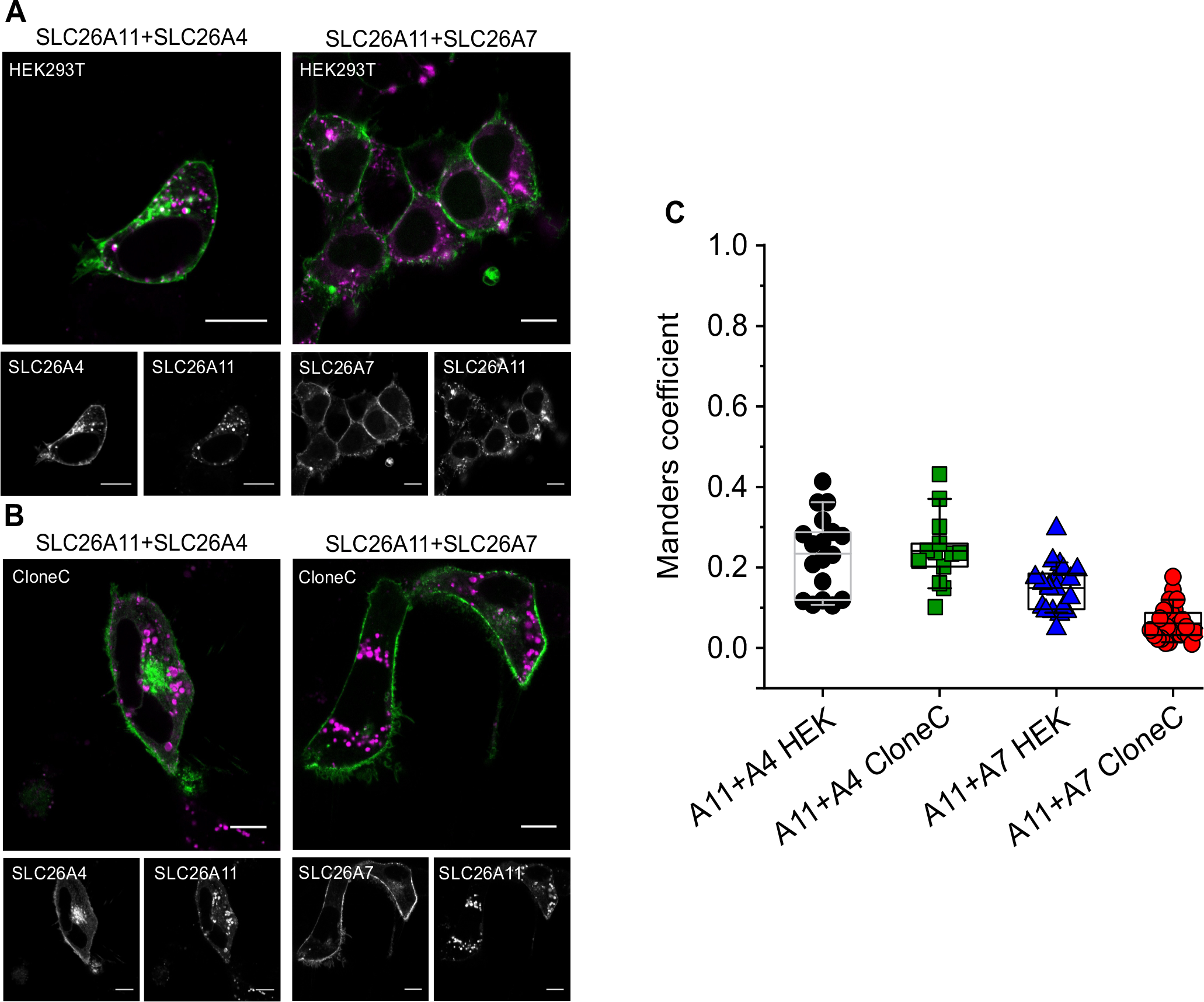
Subcellular localization of SLC26A11 after co-expression with SLC26A4/pendrin or SLC26A7. *A* and *B*, Confocal images of HEK293T cells (*A*) and ß-intercalated Clone C cells (*B*) co-expressing both indicated SLC26 isoforms. *C*, box plot of Manders coefficients of the co-localization analysis between SLC26A4/pendrin together with SLC26A11 and SLC26A7 together with SLC26A11 (SLC26A4/SLC26A11_HEK293T_, n = 18; SLC26A4/SLC26A11_CloneC_, n = 13; SLC26A7/SLC26A11_HEK293T_, n = 20; SLC26A7/SLC26A11_CloneC_, n = 25; values are given as means ± S.D. with n being the number of co-transfected cells analyzed). Scale bars = 10 µm.

### Native gel electrophoresis demonstrates exclusive homodimerization of SLC26 exchangers

We next used native gel electrophoresis (hrCNE) (Fig. 5) as an independent and well-established method to investigate membrane protein oligomerization (30-32). These experiments were combined with SDS PAGE to compare relative protein expression levels. SDS PAGE separates SLC26A4-eGFP with an apparent molecular weight of 120 kDa, closely similar to the calculated MW of 112 kDa (Fig. 5*A*). A very faint band with an apparent MW of 145 kDa above the main SLC26A4-eGFP protein band represents complex glycosylated SLC26A4-eGFP protein that can be deglycosylated by PNGaseF, but not by EndoH (Suppl. Fig. S3). SLC26A4-eGFP is expressed at similar levels in single or co-transfection experiments with SLC26A11 (Fig. 5*A*). Since the separation of homo- and heterodimers by hrCNE is based on size differences, we used an untagged version of SLC26A11. These experiments were visualized by Western blot analysis. SDS PAGE resolved two main SLC26A11 bands, one with an apparent MW of 60 kDa (calculated MW 64 kDa) for non-glycosylated and an additional with 70 kDa for complex glycosylated transporters (Suppl. Fig. S3). Co-expression with SLC26A4-eGFP (Fig. *5A*) or SLC26A7-YFP (Fig. 5*B*) resulted in slightly reduced expression level for SLC26A11, without affecting migration in SDS PAGE.

**Figure 5.**
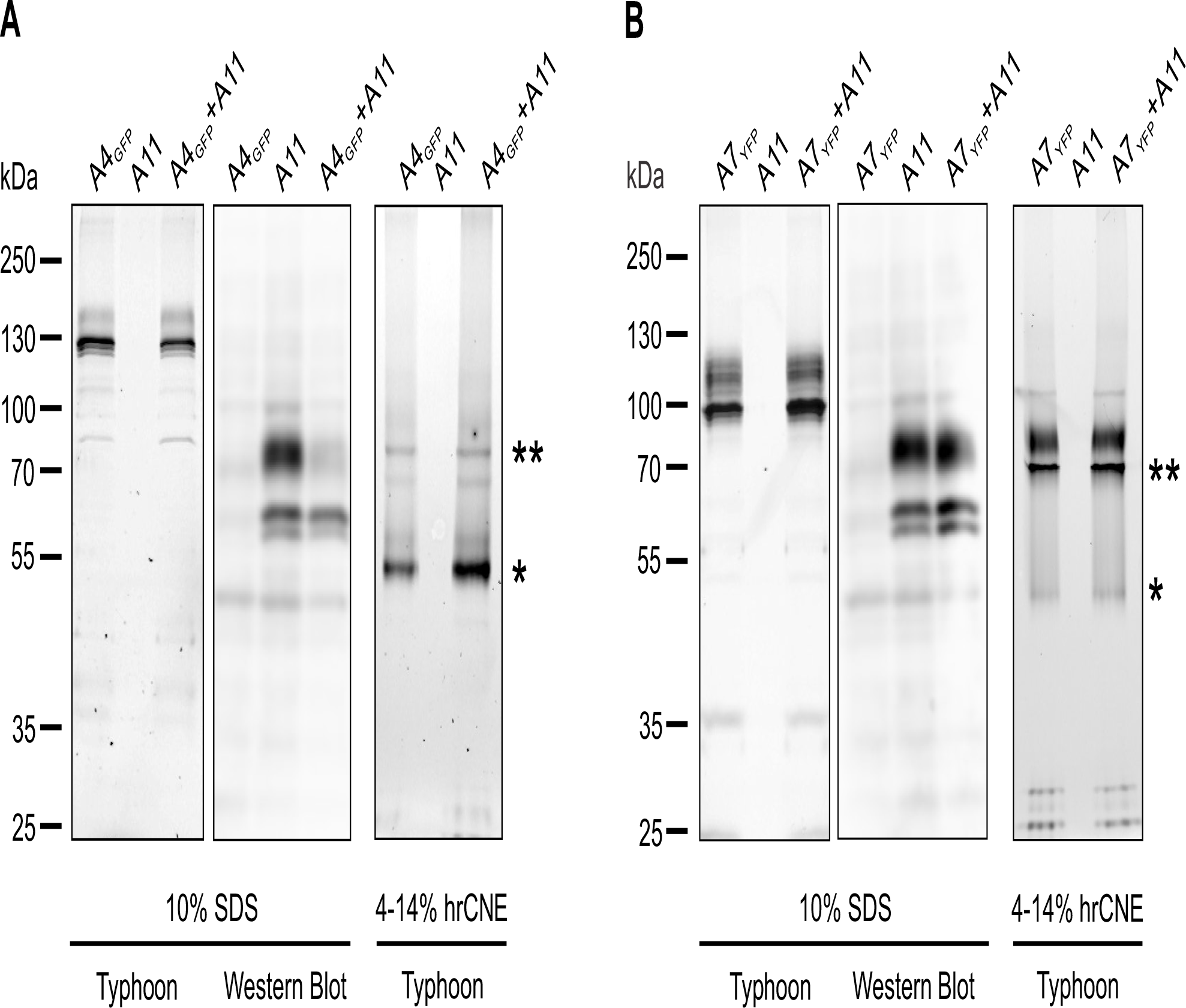
hrCNE experiments indicate no heterodimer formation of SLC26A11 with SLC26A4/pendrin or SLC26A7. *A* and *B*, SDS PAGE analysis of lysates from HEK293T cells expressing SLC26A4-eGFP, SLC26A11 or SLC26A4-eGFP together with SLC26A11 (*A*) and SLC26A7-YFP, SLC26A11 or SLC26A7-YFP together with SLC26A11 (*B*). Protein bands were visualized using fluorescent scanning (*A* or *B*, left) or Western Blot analysis (*A* or *B*, middle) using anti-SLC26A11 specific monomeric antibody (17D1). hrCNE analysis of the same cell lysates (*A* or *B*, right) indicate formation of transporter monomer (*) or transporter homodimer bands (**). Monomer as well as homodimer bands appear as double bands due to its glycosylation status, non-glycosylated protein as lower band and glycosylated protein as upper band.

When expressed alone, hrCNE analysis of SLC26A4-eGFP revealed two distinct bands (Fig. 5A, right), a predominantly monomeric band and a faint homodimeric band (lower and upper black stars), indicating that SLC26A4-eGFP forms more stable monomers than homodimers. In comparison to that, hrCNE of SLC26A11-eGFP or SLC26A11-mCherry shows two high molecular weight protein bands and one faint low molecular weight protein band, indicating the formation of more stable homodimers than monomers. SLC26A11 can be complex glycosylated, which causes the appearance of protein double bands (Suppl. Fig. S4, lower and upper black stars). Since untagged SLC26A11 exhibit a lower molecular mass than SLC26A4-eGFP, SLC26A4-SLC26A11 heterodimerization should give rise to a new protein band migrating between the SLC26A4-eGFP dimer and monomer band (33). Co-transfection of SLC26A4 and SLC26A11 did not result in novel fluorescent bands, providing additional evidence against the formation of heterodimeric assemblies.

Figure 5*B* illustrates the comparable analysis for SLC26A7-YFP and SLC26A11. SLC26A7-YFP exhibits an apparent molecular weight of 100 kDa (calculated MW 98 kDa), with complex glycosylation increasing the apparent MW to 120 kDa (Suppl. Fig. S3). SLC26A7-YFP expression levels were unaffected by co-transfection with SLC26A11 (Fig. 5*B*, left), and Western blot analysis provided similar expression levels of SLC26A11 expressed alone or co-transfected together with SLC26A7-YFP (Fig. 5*B*, middle). hrCNE resolved two high molecular SLC26A7-YFP bands, when expressed alone, corresponding to glycosylated and non-glycosylated homodimers (Fig. 5*B*, right, lower and upper black stars), in addition to one faint monomeric protein band. SLC26A7-YFP forms more stable homodimers than monomers. In co-expression experiments of SLC26A7-YFP with untagged SLC26A11, the same fluorescent bands were observed, indicating that SLC26A7-YFP and SLC26A11 do not co-assemble into heterodimers.

### Co-expression of SLC26A11 with SLC26A1, SLC26A2, SLC26A6, and SLC26A9 leaves its lysosomal localization unaffected

Since SLC26A11 is the only SLC26 isoform with predominant intracellular localization (Fig. 1), we used co-expression in HEK293T cells and confocal imaging as test for co-assembly of SLC26A11 with other SLC26 isoforms, such as SLC26A1, SLC26A2, SLC26A6, and SLC26A9™. Expression of SLC26A3 was reduced to levels below the detection limit upon coexpression with SLC26A11, preventing any analysis for this combination. Since SLC26A5/prestin is exclusively expressed in inner ear outer hair cells, we refrained from studying oligomerization between SLC26A11 and this SLC26 family member.

Figure 6 shows representative confocal images from HEK293T cells co-expressing SLC26A11 with SLC26A1 (Fig. 6*A*), SLC26A2 (Fig. 6*B*), SLC26A6 (Fig. 6*C*) and SLC26A9™ (Fig. 6*D*). Since cells expressing SLC26A9-eGFP show substantial intracellular staining that might make the identification of SLC26A11 heterodimers with separate subcellular distribution difficult; we decided to co-express the truncation mutant SLC26A9™ with improved surface membrane insertion (Fig. 1). Manders coefficients summarizing the subcellular unchanged subcellular localizations for SLC26A11 or the co-transfected SLC26 isoform for all tested combinations (Manders coefficient (MC)_SLC26A1/SLC26A11_ = 0.12 ± 0.05 (n=18), (MC)_SLC26A2/SLC26A11_ = 0.09 ± 0.03 (n=24), (MC)_SLC26A6/SLC26A11_ = 0.09 ± 0.04 (n=24), and (MC)_SLC26A9TM/SLC26A11_ = 0.22 ± 0.11 (n=24) (Fig. 6*E*). We conclude that SLC26A11 does not form heterodimeric assemblies with other SLC26 isoforms.

**Figure 6.**
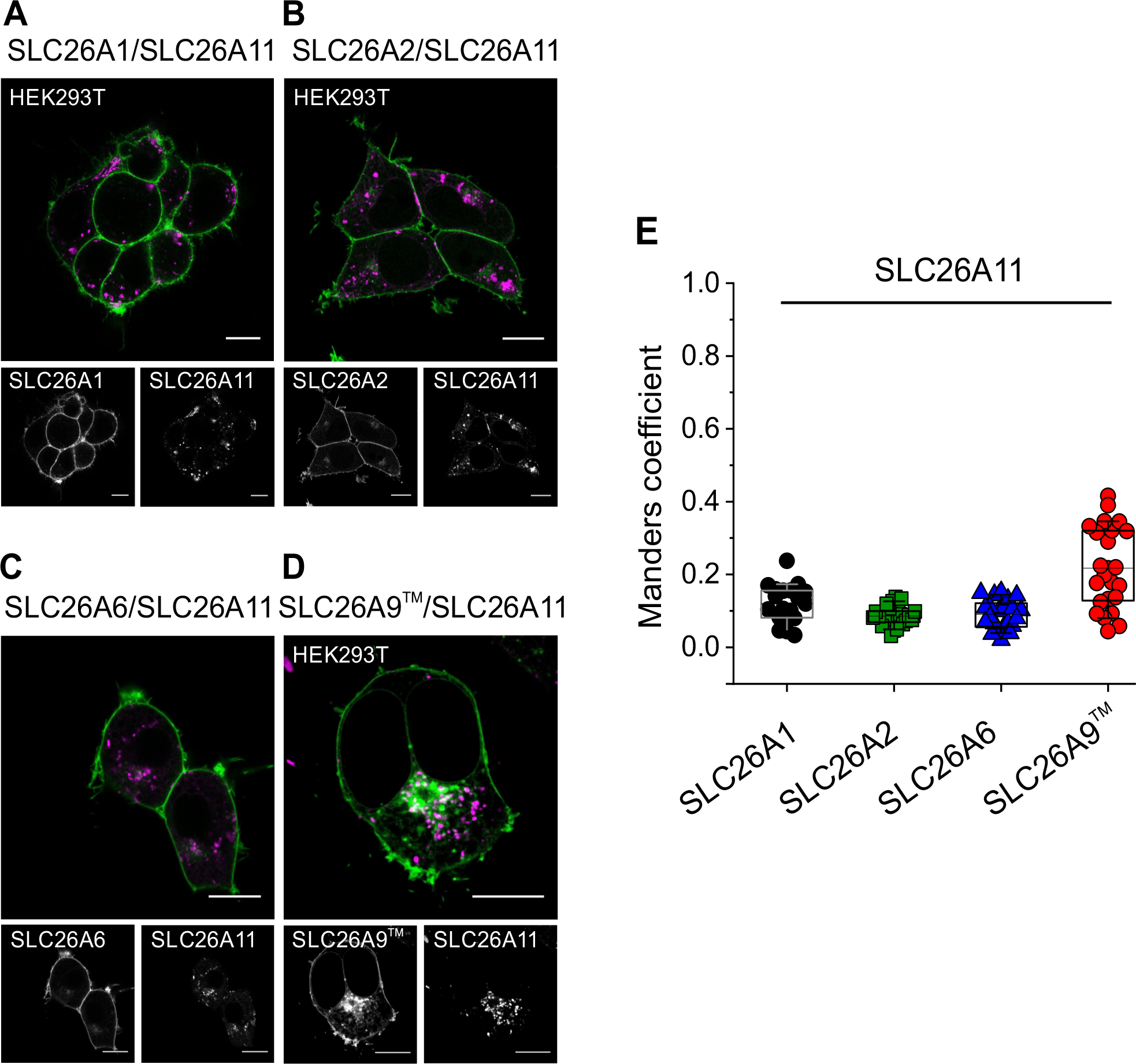
Comparison of subcellular localization of SLC26A11 heterologously co-expressed in HEK293T cells with SLC26A1, SLC26A2, SLC26A6 or SLC26A9™. *A - D*, Confocal images of HEK293T cells co-expressing SLC26A11 together with SLC26A1 (*A*), SLC26A2 (*B*), SLC26A6 (*C*) or SLC26A9™ (*D*). *E*, box plot of Manders coefficients of co-localization analysis between SLC26A11 and the corresponding SLC26 transporter (SLC26A11/SLC26A1, n = 18; SLC26A11/SLC26A2, n = 24; SLC26A11/SLC26A6, n = 24; SLC26A11/SLC26A9™, n = 24; values are given as means ± S.D. with n being the number of co-transfected cells analyzed). Scale bars = 10 µm.

## Discussion

Thus far, no agreement has been reached about transport functions, interaction partners, or subcellular distributions of SLC26A11, making this isoform one of the most enigmatic members of the SLC26 family. Whereas initial characterization suggested SLC26A11 functioning as electroneutral sulfate transporter (13,14), later work reported chloride currents associated with SLC26A11 (15). These findings, together with altered intracellular [Cl^-^] in Purkinje neurons from brain-specific knock-out animal (16) suggested SLC26A11 functioning as anion channel. SLC26A11 is broadly expressed in many cell types (14), but, thus far, the consequences of constitutive SLC26A11 ablation in all cell types remain unknown. In contrast to other SLC26, *SLC26A11* sequence variants have not yet been linked to any human genetic disease. Thus, neither the molecular function nor the cellular roles of SLC26A11 are sufficiently understood.

We here demonstrate that SlC26A11 differs from other SLC isoforms in its subcellular distribution (Fig. 1). While SLC26A1, SLC26A2, SLC26A3, SLC26A4/pendrin, SLC26A5/prestin, SLC26A6, SLC26A7, and SLC26A9 show predominant surface membrane insertion in heterologous expression experiments with FP-fusion proteins, we observed exclusive localization of SLC26A11 to late endosomal compartments, without visible or measurable surface membrane insertion (Fig. 1). The predominant lysosomal localization is in agreement with proteomics data (34), but in disagreement with a general consensus reached by physiologists over recent years. After heterologous expression in *Xenopus* oocytes, predominant surface membrane localization of SLC26A11 was observed by confocal immunofluorescence microscopy experiments (13). Moreover, anion currents could be measured in microelectrode voltage clamp experiments (15). There might be differences between mammalian cells and amphibian oocytes. Moreover, overexpression in heterologous systems might result in findings that differ from native cells, and mammalian cell lines and oocytes might produce different levels of overexpression. We tried to test for such influences using different expression vectors (Suppl. Fig. S1), without discernable influence. Another result that supports localization in the surface membrane is the reduced intracellular [Cl^-^] in neurons after genetic ablation of SLC26A11 (16). Clearly, as consequence, the subcellular localization of SLC26A11 will have to be tested in native cells/ tissues.

Hetero-oligomerization can increase the functional diversity in a protein family. In ion channels, hetero-oligomerization modifies channel gating and ion conduction as well as the pharmacological properties, but also the subcellular distribution (18,19,21,35-37). For transporters, functional intersubunit interactions are difficult to study, and thus far, only hetero-oligomerization-associated changes in subcellular distribution have been reported (29,33,38). Homo- and hetero-dimerization has been extensively studied for CLC anion transporters (39). There exist five intracellular CLC-type Cl^-^ H^+^ exchangers (ClC-3, ClC-4, ClC-5, ClC-6 and ClC-7) that differ in functional properties as well as in subcellular localizations (39,40). Among those, only ClC-3 forms heterodimers with ClC-4 and permits ClC-4 targeting to various compartments of the endo/lysosomal systems as heterodimer (29,41,42). Exclusive homodimerization restricts the subcellular localization of the other CLC transporters and permits targeting of transporters with certain functional properties to selected organelles. For example, exclusive homodimerization of the lysosomal CLC-7 prevents targeting to other intracellular localizations as part of a heterodimeric assembly. SLC26 anion exchangers form dimeric assemblies that can be demonstrated by structural studies (22,43-48) as well as by native gel electrophoresis (49). Since both experimental approaches do not exclude formation of higher oligomers, which are not sufficiently stable to tolerate solubilization/purification, we recently used a new dual-color colocalization (DCC) approach for the accurate determination of subunit stoichiometries of membrane proteins *in situ* (50,51). This approach also demonstrated that a dimeric assembly is the predominant subunit stoichiometry of SLC26 proteins, without indications of higher oligomeric states.

We here studied whether SLC26A11 hetero-multimerization with other SLC26 isoforms might result in novel subcellular distributions. Since SLC26A11 resides in lysosomes and expression of all other tested SLC26 isoforms result in predominant plasmalemmal localizations (Fig. 1), heterodimer formation must result in a change of localizations, i.e. of SLC26A11 into cell compartments other than the lysosome and/or of other SLC26s from the plasma membrane to intracellular compartments. This was not observed, providing strong evidence against heterodimerization of these isoforms. This analysis is semi-quantitative: although we cannot exclude minor fractions of heterodimers, it demonstrates a vast majority of SLC26A11 remaining in homodimeric assemblies when coexpressed with other isoforms. Confocal imaging was performed in living cells, and thus all types of interactions - that might modify the localizations of the tested proteins - can be assessed. It is not affected by experimental modifications as for example detergent stability of complexes. Since we use fluorescent fusion proteins, added tags might affect associations with other proteins. However, since we use comparable linking strategies and since the tags do not affect homodimerization, this possibility appears unlikely.

Native gel electrophoresis has been extensively used for studying membrane protein oligomerization (31,49,52-54). It relies on solubilization steps that might disrupt complexes that exist in living cells. Different oligomeric states are distinguished by size differences, and we therefore expressed SLC26A4/pendrin or SLC26A7 GFP fusion proteins together with untagged SLC26A11. We did not observe fluorescent proteins that differed from SLC26A4/SLC26A7 monomers or homodimers in co-expression experiments with SLC26A11, providing additional evidence against the formation of heterodimers between SLC26A11 with these two isoforms.

Our results support a lysosomal localization of homodimeric SLC26A11, with no evidence of heterodimer formation that would lead to distinct subcellular localization. SLC26A11 functioning as pH-dependent Cl^-^/SO_4_^2-^ exchanger and/or as passive Cl^-^ conductor (15) might contribute to lysosomal ion homeostasis. Since lysosomes metabolize sulfur in amino acids, lipids and sugars to SO_4_^2-^, effective SO_4_^2-^ export is necessary for normal lysosome function, possibly assigning an important housekeeping function to SLC26A11. Functional studies under ionic conditions resembling those found in lysosomes will facilitate a better understanding of SLC26A11’s transport functions and its contribution to normal cellular processes.

### Experimental procedures

#### Plasmid construction

cDNA encoding full length SLC26A1 and SLC26A2 (kindly provided by Dr. Tomohiro Shima, Tokyo, Japan) were sub-cloned into pEGFP-N1 vector (Clontech, Germany) or pcDNA5/FRT/TO vector (Novagen, Germany). cDNAs encoding full length WT SLC26A3, SLC26A6, SLC26A9 (kindly provided by Dr. David Mount, Boston USA), SLC26A5/prestin (kindly provided Dr. Bernd Fakler, Freiburg, Germany) and SLC26A7 (kindly provided by Dr. Karl Kunzelmann, Regensburg, Germany) were sub-cloned into pcDNA5 FRT/TO vector (Novagene, Germany). Truncated SLC26A9™ was generated via overlapping PCRs according to (22). cDNA encoding full length human SLC26A4/pendrin was kindly provided by Dr. Dominik Oliver sub-cloned into pEGFP-N1 vector. cDNA encoding full-length mouse SLC26A11 (kindly provided by Dr. Rainer Schreiber, Regensburg, Germany) cloned into pTarget vector (Promega, Germany) was subcloned into FsY1.1 G.W. vector (kindly provided by Dr. Mikhail Filippov, Nizhny Novgorod, Russia), pcDNA3.1.(+) or pcDNA5 FRT/TO vector (Novagene, Germany). Human SLC26A11 cDNA (kindly provided by Dr. Erik Geertsma, Dresden) was sub-cloned into pcDNA3.1 vector (Novagen, Germany). Enhanced green, monomeric cherry or monomeric yellow fluorescent proteins (eGFP, mCherry or mYFP) were fused in frame to the C-Terminus of the proteins, respectively). For biochemical analysis, untagged SLC26A11 - that was not fused to the coding sequence of a fluorescent protein - was analyzed.

#### Confocal Microscopy and Image Analysis

For confocal imaging of living cells, HEK293T cells or CloneC cells (ATCC ^®^ CRL2531 ™) were either transfected with SLC26 fluorescent fusion proteins alone or in combination with plasmids encoding fluorescent Lamp1-mCherry or fluorescent Calnexin-mCherry using Lipofectamine 2000 (Invitrogen). We received the lysosomal marker Lamp1 as a gift from Walter Mothes, Addgene plasmid #1817 (55), and the ER marker Calnexin as gift from Michael Davidson (http://n2t.net/addgene:55005; RRID: Addgene_55005). Cells were plated on poly-L-lysine-coated coverslips 6 h after transfection, and images were taken 48 h later with a Leica TCS SP5 II inverted microscope (Leica Microsystems, Wetzlar, Germany) using a 63 x /1.40 NA oil immersion objective in phosphate-buffered saline (PBS; 130 mM NaCl, 7 mM Na_2_HPO_4_, and 3 mM NaH_2_PO_4_; pH: 7.4) at room temperature. Images were digitalized with a resolution of 1024 x 1024 pixels, 400 Hz velocity, and 5-line average in sequential scanning mode. eGFP and mYFP were excited with a 488-nm Ar-laser and mCherry with a 594-nm He-Ne-laser. Emission signals were detected after filtering with a 500-550 or 600-650 nm bandpass filter. Confocal images were processed for publications using ImageJ 1.44p software (Image J v.1.53c, Wayne Rasband, National Institutes of health, Bethwsda, Rockville, MD, United States) (56).

### Generation of anti-SLC26A11 antibody, 17D1

We expressed a fusion protein of the STAS domain of murine SLC26A11 (amino acids 451-593 with the sequence: varpktqvse gqifvlqpas glyfpaidal reaitnrale aspprsavle cthissvdyt vivglgelle dfqkkgvala fvglqvpvlr tllaadlkgf ryfttleeae kflqqepgte pnsihedavp eqrssllksp sgp) with a carboxy-terminal deca-His-Tag for affinity purification. The amino acid sequence differs from other SLC26 protein family members and is therefore suitable to reduce potential antibody cross detection. Rats (strain: Lou/C) were immunized with the purified protein using standard procedures (R. Feederle, Antibody Core Facility, Helmholtz Zentrum München), and spleen cells were fused to a mouse myeloma cell line (P3X63-Ag8.653) to generate hybridoma cells. The primary supernatant was tested by ELISA and Westernblot analysis. Hybridoma cells were sub-cloned via single-cell distribution until all reacted positively. The final clone was expanded, and used to generate supernatant containing the monoclonal antibody (17D1; subtype IgG_2_). To test the antibody, HEK293T or Clone C cells were transfected with either SLC26A11-eGFP fusion construct or non-tagged SLC26A11 construct and plated on poly-L-lysine-coated coverslips 6 h after transfection. 48 hours later, cells were fixed for 5 minutes in 4% paraformaldehyde, washed with phosphate-buffered saline and incubated for one hour with primary antibody diluted in 5% Chemiblocker (Chemicon), 0.5% Triton X-100, and 0.05% NaN_3_ in phosphate-buffer. Monoclonal 17D1 antibody (rat IgG_2_ subtype) was detected and visualized with donkey-anti-rat Cy3 in cells expressing SLC26A11-eGFP and with don-key-anti-rat Alexa488 in cells expressing non-tagged SLC26A11. Cells were examined using a confocal laser scanning microscope (Leica TCS SP5 II inverted microscope (Leica Microsystems, Wetzlar, Germany) using a 63 x /1.40 NA oil immersion objective. Confocal images were processed for publications using ImageJ 1.44p software (Image J v.1.53c, Wayne Rasband, National Institutes of health, Bethwsda, Rockville, MD, United States) (56). Cross detection of SLC26A4/pendrin, SLC26A7, and SLC26A9 was tested via Western Blot analysis using 1D17 (1:10 dilution) on whole cell lysates (Supplementary Fig. 5). Approximately 10 μg whole-cell lysate sample per lane was separated by 10% SDS PAGE and no cross detection was found. Western blot protocol see below.

### Biochemical Analysis

For biochemical analyses, SLC26A4-eGFP, SLC26A7-YFP or untagged SLC26A11 were transiently expressed in HEK293T cells alone or HEK293T cells co-transfected with plasmids encoding either for SLC26A4-eGFP together with SLC26A11 or for SLC26A7-YFP together with SLC26A11. 24 hours after transfection, HEK293T cells were washed with ice cold PBS and lysed with buffer containing 0.1 M sodium phosphate, pH 8.0, 0.5% digitonin, a protease inhibitor cocktail and 20 mM iodoacetamide as previously described (29,57). An aliquot of approximately 10 µg of generated whole-cell-lysate was analyzed by 10% SDS PAGE. Gels were either scanned using a fluorescence gel scanner (see below), or proteins were transferred for Western blot analysis onto a polyvinylidene fluoride membrane (PVDF, Merck/Millipore, Germany). Nonspecific binding sites were blocked with 1% (w/v) dry milk in PBS supplemented with 0.05% (v/v) Tween-20 (PBS-T) for 30 minutes. The membrane was incubated with primary 17D1 antibody (1:10 dilution) in PBS-T for one hour at room temperature. After rinsing the membrane three times with PBS-T for 5 min each, secondary antibodies conjugated to horseradish peroxidase (donkey anti-rat-HRP; dilution, 1:5000 (Sigma-Aldrich/Merck, Germany)) in PBS-T were added for 45 minutes at room temperature. After rinsing the membrane three times with PBS-T for 5 min each and one time with PBS for 5 min, signals were visualized with an enhanced chemiluminescence detection system (SuperSignalTM West Pico PLUS, ThermoFisher, Germany). Signals were visualized using a GeneGnome (Synoptics, UK). For high-resolution clear native gel electrophoresis (hrCNE), native 4-14% acrylamide-gradient gels were casted as described previously (30,31). The anode buffer contained 25 mM imidazole/HCl, pH 7.0. The cathode buffer contained 50 mM tricine, 7.5 mM imidazole, pH 7.0, 0.05% sodium-deoxycholate, and 0.01% DDM (30). Approximately 10 μg whole-cell lysate samples were run at 8°C for a total of 3 h (100 V for 1 h, followed by 150 V for 2 h).

Gels were scanned using a fluorescence gel scanner (Typhoon FLA9500, GE Healthcare, Freiburg, Germany) at 100 μm resolution. eGFP and mYFP were excited at 473 nm, and its emissions recorded using a 530/20 bandpass filter. Gel images were visualized using ImageJ 1.44p software. Gels were shown in black and white and the appearance of the entire gel was adjusted using the Brightness and Contrast tool of ImageJ.

## Supporting information

Supplemental Figures Bungertetal_2024

## Data availability

The original contributions presented in the study are included in the article/Supplementary Material, further inquiries can be directed to the corresponding author.

## Supporting information

This article contains supporting information.

## Acknowledgements

The coding sequences for SLC26 transporters were kindly provided by Dr. Tomohiro Shima, Tokyo, Japan, Dr. David Mount, Boston USA, Dr. Bernd Fakler, Freiburg, Germany, Dr. Karl Kunzelmann, Regensburg, Germany, Dr. Dominik Oliver, Marburg, Germany and Dr. Rainer Schreiber, Regensburg, Germany. We thank DR. Bassam Haddad and the DFG research unit FOR5046 for helpful discussions and Joachim Schmitz of excellent technical support.

## Author contributions

SBP, RG and ChF designed the study and wrote the paper. SBP performed confocal microscopy and biochemical analyses. All authors approved the final version of the manuscript.

## Funding and additional information

These studies were supported by the Deutsche Forschungsgemeinschaft (DFG, German Research Foundation) to Ch.F. (FA 301/15-1) as part of the Research Unit FOR5046

