## Supplemental Figures Bungertetal_2024 for "Oligomerization and cellular localization of SLC26A11"

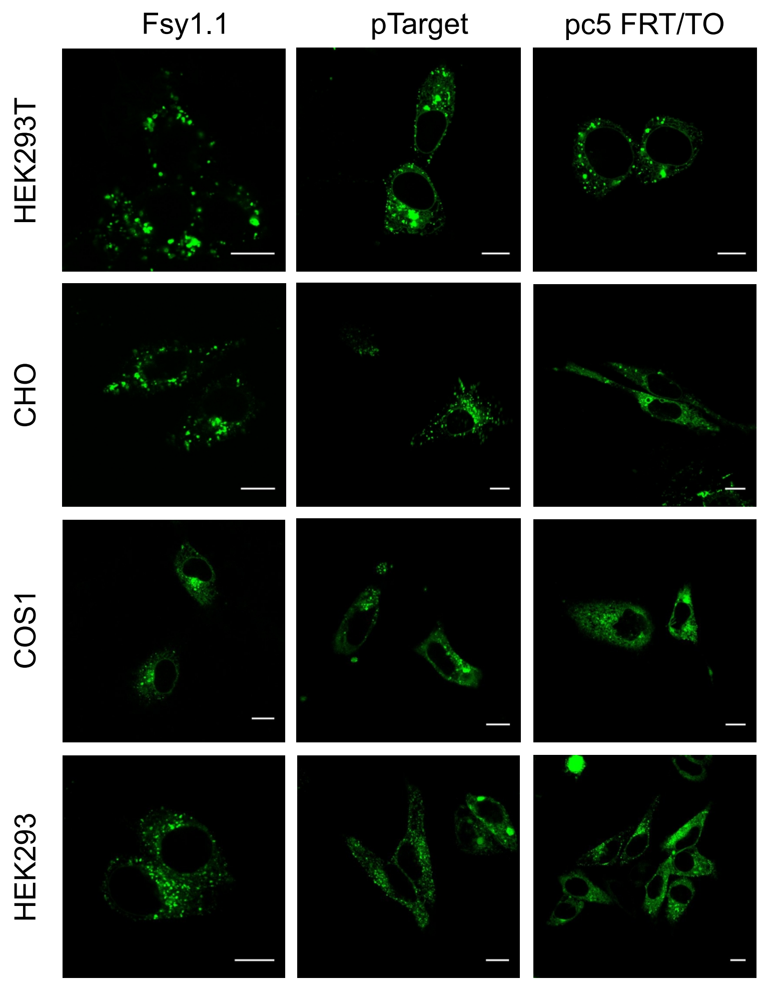


**Supplementary Fig. 1. Expression of SLC26A11 in different cell lines using different expression plasmids.** Confocal images of HEK293T, CHO, COS1 and HEK293 cells heterologously expressing SLC26A11-eGFP using the expression plasmids Fsy1.1 GW, pTarget or pcDNA5 FRT/TO. SLC26A11 is located intracellular. Scale bars 10 µm.

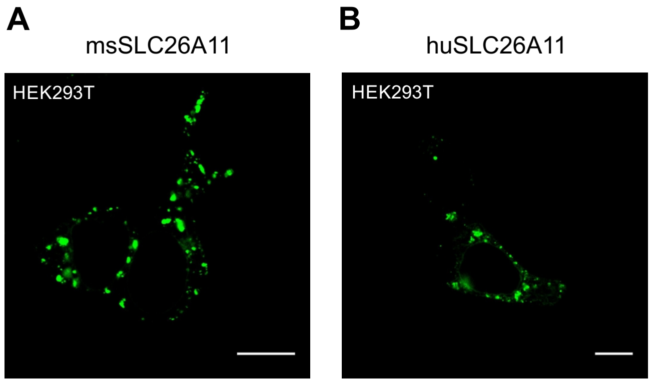


**Supplementary Fig. 2 Subcellular localization of mouse SLC26A11 and human SLC26A11 protein in HEK293T cells.** A and B, Confocal images of HEK293T cells heterologously expressing mSLC26A11-eGFP (A) or hSLC26A11-eGFP (B). Scale bars 10 µm.

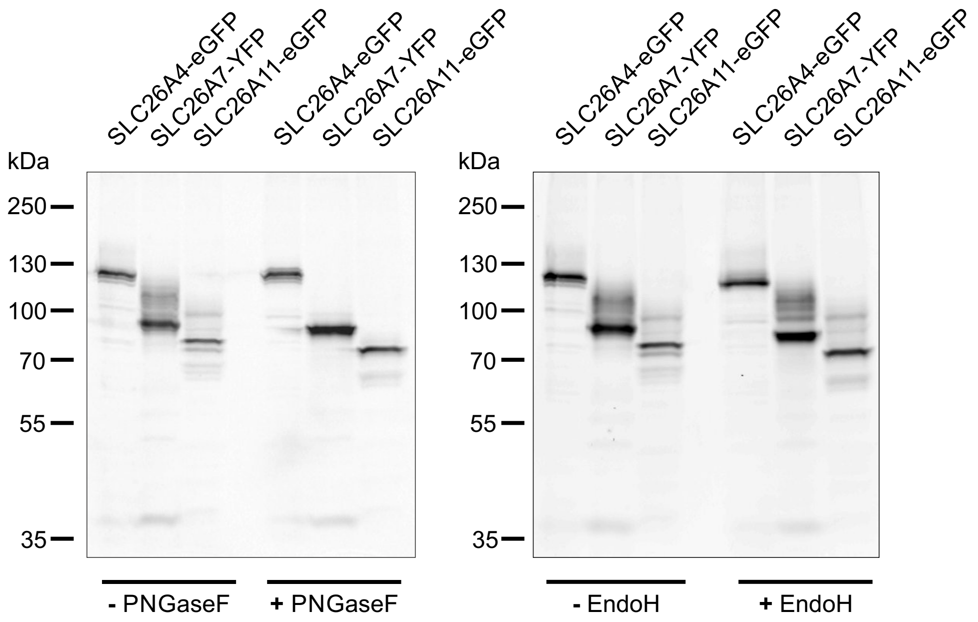


**Supplementary Fig. 3 Glycosylation analysis of SLC26A4/pendrin, SLC26A7 and SLC26A11 proteins.** Representative SDS PAGE of lysates from HEK293T cells expressing SLC26A4-eGFP, SLC26A7-YFP or SLC26A11-eGFP before and after PNGaseF treatment (left) or EndoH treatment (right).

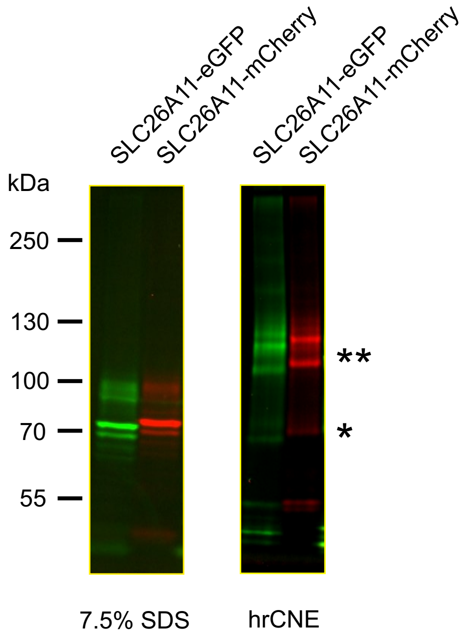


**Supplementary Fig. 4 Comparions of SDS PAGE and hrCNE analysis of lysates from HEK293T cell expressing SLC26A11-eGFP and SLC26A11-mCherry.** SDS PAGE shows SLC26A11-eGFP and SLC26A11-mCherry with an apparent MW of 75 kDa (calculated MW 91 kDa) for non-glycosylated and higher MW bands of 90 kDa for complex glycosylated transporters (left). hrCNE indicates two pronounced high molecular bands corresponding to glycosylated and non-glycosylated homodimers (**) and a faint monomer band (*) (right).

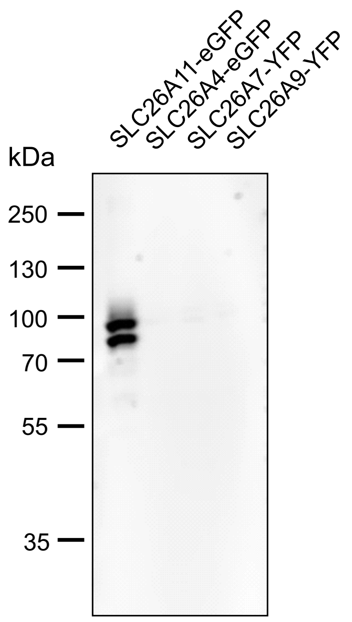


**Supplementary Fig. 5 Cross detection of SLC26A4, SLC26A7, or SLC26A9 with the monoclonal antibody rt17D1.** 10% SDS PAGE of whole cell lysates from HEK293T cells expressing SLC26A4/pendrin, SLC26A7, SLC26A9, or SLC26A11 and Western Blot analysis using rt1D17 monoclonal antibody (1:10 dilution). Approximately 10µg whole-cell lysate per lane was loaded and no cross detection was found.
